# Disturbance macroecology: integrating disturbance ecology and macroecology with different-age post-fire stands of a closed-cone pine forest

**DOI:** 10.1101/309419

**Authors:** Erica A. Newman, Mark Q. Wilber, Karen E. Kopper, Max A. Moritz, Donald A. Falk, Don McKenzie, John Harte

## Abstract

Macroecological studies have generally restricted their scope to relatively steady-state systems, and as a result, how biodiversity and abundance metrics are expected to scale in disturbance-dependent ecosystems is unknown. We examine macroecological patterns in a fire-dependent forest of Bishop pine (*Pinus muricata*). We target two different-aged stands in a stand-replacing fire regime, one a characteristically mature stand with a diverse understory, and one more recently disturbed by a stand-replacing fire (17 years prior to measurement). We compare the stands using macroecological metrics of species richness, abundance and spatial distributions that are predicted by the Maximum Entropy Theory of Ecology (METE), an information-entropy based theory that has proven highly successful in predicting macroecological metrics across a wide variety of systems and taxa. Ecological patterns in the mature stand more closely match METE predictions than do data from the recently disturbed stand. This suggests METE’s predictions are more robust in late-successional, slowly changing, or steady-state systems than those in rapid flux with respect to species composition, abundances, and organisms’ sizes. Our findings highlight the need for a macroecological theory that incorporates natural disturbance and other ecological perturbations into its predictive capabilities, because most natural systems are not in a steady state.

## Introduction

Disturbance is pervasive in ecosystems, and it influences patterns of species diversity, abundance, and community membership over space and through time (Turner 1989). However, macroecology, the discipline concerned with large-scale patterns of diversity, has generally avoided studies of disturbed systems for at least two reasons: first, that disturbed systems are perceived as being in “transition” and unlikely to produce replicable, generalizable results; and second, assumptions of steady-state, equilibrium and stabilizing mechanisms in macroecological theory are common and often required in order to solve equations (see Hubbell 2001). Macroecology has instead primarily focused on ecosystems that are perceived to be relatively stable (Fisher et al. 2010), in that they exhibit low variance in community structure through time (Turner et al. 1993). Disturbed ecosystems (and patches within ecosystems) are likely to be in flux with respect to species composition and richness, species-area relationships, distribution of abundances, and body sizes, and intraspecific spatial distributions of individuals. This is true of ecosystems that have recently undergone, or are continuing to undergo, natural disturbances (those that are part of a repetitive disturbance regime, *sensu* Turner 2010), anthropogenic changes, and other ecological disruptions. The dynamics of disturbed sites and entire disturbance regimes are not captured by standard macroecological study systems, which are often chosen because they are in or near steady states (e.g. most of the Center for Tropical Forest Science plots represent late-successional, primary forest; Condit 1998 Ch.1).

Although natural disturbances have both large- and small-scale structuring effects in all ecosystems (Turner 1989), no macroecological study, to our knowledge, has addressed how metrics of biodiversity and abundance scale in disturbance-dependent ecosystems. Various studies on succession have led macroecologists to invoke “disturbance” broadly (including human activities, environmental variability, invasive species and so on) as a factor responsible for deviations from theoretical predictions or expected patterns, although it remains unclear whether macroecological patterns reported across ecosystems are properties of undisturbed, steady-state communities or are properties of all ecological systems. This failure to incorporate disturbance into macroecology poses a major challenge to the utility of this field in understanding ecological dynamics as well as global change. Synthesizing a “macroecology of disturbance” that incorporates quantitative macroecological metrics could have considerable consequences for conservation efforts, given that many ecosystems with active disturbance regimes (and the species that have evolved in them) rank among the most globally endangered (Turner 2010; and see Noss et al. 1995, Schlossberg and King 2015, Batllori et al. 2013). Distinguishing the effects of natural disturbances from those of anthropogenic changes is also important for predicting future states of ecosystems.

Here, we will restrict the use of the term “disturbance” to refer to “natural disturbances,” which satisfy the following four characteristics: a) they cause mortality of individual organisms in a community; b) however, they do not cause mortality of all individuals in the community and therefore do not result exclusively in primary succession; c) they are part of a historic and repetitive “disturbance regime” (Turner 2010) with well-defined characteristics (Pickett and White 1985, Turner et al. 1998, Turner 2010); and d) the disturbance is “absolute” rather than “relative” (Pickett and White 1985, White and Jentsch 2001) in that each disturbance event is “a relatively discrete event in time that disrupts the ecosystem, community or population structure and changes the resources, substrate availability or physical environment” (White and Jentsch 2001) (Appendix A). This strict operational definition of “disturbance” as synonymous with “natural disturbance” is consistent with its usage in several influential reviews of disturbance ecology (Pickett and White 1985, White and Jentsch 2001, Turner 2010).

Past macroecological work that incorporates ecological disturbances of any type focused predominantly on their effects on the shape of the species-abundance distribution (SAD). Although the SAD is well-studied (reviewed in McGill et al. 2007; White et al. 2012; Baldridge et al. 2015), the underlying shape of a “natural” or generic SAD is debated (see for example Hill et al. 1995, Nummelin 1998, Hill and Hamer 1998; Ulrich et al. 2010), and various distributions have been proposed. Empirical support for each of these distributions is mixed. For the rank-abundance form of the SAD, a lognormal distribution is reported from many steady-state systems (Whittaker 1965; May 1975; Gray 1981; Ulrich et al. 2010), while other studies (Dennis and Patil 1979, Kempton and Taylor 1974) including “big data” approaches showing that the log series distribution may be the most common across systems and taxa (White et al. 2012; Baldridge et al. 2015). One study suggests the prevalence of the “double geometric” distribution (Alroy 2015). One-time ecological perturbations are often invoked as responsible for a lognormal SAD (Bazzaz 1975; Hill and Hamer 1998; Kempton and Taylor 1974; Death 1996; Newman et al. 2014). Kempton and Taylor (1974) show in a comparative study that moth communities in undisturbed plots sites in the Rothamsted Insect Survey in England exhibit log series SADs, and plots recovering from agricultural activity exhibit lognormal SAD. Certain ecological factors, sampling methods (Ulrich et al. 2010), detection issues (Tokeshi 1993), and mathematical processes (such as the central limit theorem) may also produce the lognormal (Tokeshi 1993). Work focusing on succession suggests a transition in the shape of the SAD from geometric in early successional stages to lognormal and subsequently log series in later stages (Whittaker 1975; Bazzaz 1975). Other macroecological metrics are much less well studied in the context of ecological disruption, although the species-area relationship (SAR) has been examined through experimental work with removal of seed predators (Supp et al. 2012), and the effects of ecological disruptions and perturbations are beginning to be investigated more broadly (Supp et al. 2014, Mayor et al. 2015).

The Maximum Information Entropy Theory of Ecology (METE) is a macroecological theory (Harte et al., 2008; 2009; Harte 2011; Harte and Newman 2014) that provides a statistical framework for linking metrics that were previously considered in isolation, the SAR, the SAD, and species-level spatial abundance distributions (SSADs), a metric quantifying the spatial distribution of individuals in a species over a given area. METE relies on the maximum information entropy inference procedure (MaxEnt) to predict least-biased probability distributions, given empirical constraints (Jaynes 1982), but invokes no explicit physical or ecological mechanisms (Harte 2011, Harte and Newman 2014). An application of the MaxEnt procedure, the “ASNE” version of METE (Harte and Newman 2014) uses only the relationships between four non-adjustable state variables that take on values from the system being measured: *S*_0_ (total species), *A*_0_ (total area under consideration), *N*_0_ (total abundance), and *E*_0_ (total metabolic energy). The state variables are static, not dynamic in this formulation, and there are no adjustable parameters characterizing the scaling of species diversity, abundances, and energetics in a system. Mathematical forms of empirical constraints arise from ratios of the state variables. More complete mathematical constructions of distributions are available in Harte (2011).

Empirical tests of METE strongly support its core predictions, including the SAR and SAD (Harte et al., 2008; 2009; White et al. 2012), species-level spatial abundance distributions, and certain metabolic predictions (Newman et al. 2014, Xiao et al. 2014), but certain spatial distribution and metabolic predictions are not supported (see McGlinn et al. 2015, Newman et al. 2014, Xiao et al. 2014). METE has accurately predicted the SAR, SAD and SSAD for a range of natural communities spanning different taxa and biomes, including herbaceous plants, trees, vertebrates and invertebrates, and in temperate, tropical, and montane environments, as well as isolated island communities (Harte et al. 2008; Harte et al. 2009; Harte 2011; Rominger et al. 2016). This study represents the first assessment of these common macroecological metrics for a plant community in a high-intensity natural disturbance regime.

### Applying METE to ecosystems in transition

As applied here, METE might accurately capture “snapshots” of rapidly changing ecosystems at an instant in time, and the predictions of the ASNE version of METE are also static in time. A dynamic version, while desirable (Fisher et al. 2010), is not yet available. Here, separate plots are placed to capture the macroecological patterns that characterize live, aboveground plant communities within separate patches in a disturbed landscape. As is the case for many macroecological studies, we here choose to study living, aboveground plants above a certain size threshold, which are a high-detectability system, with relatively large, sessile organisms. “Propagules” or organisms that persist through major disturbances in some form (as in a seed bank) are essential to a complete understanding of the disturbance ecology of a particular system, and may make up the larger part of plant abundances in an area. These propagules are not included in our measurements, but METE predictions hold for the remaining community of interest, and metrics scale with the measured community, rather than the full community (Harte et al. 2013). METE makes the least biased predictions of the community of interest as characterized by the measured state variables (Harte 2011) as a result of its underlying

MaxEnt formalism (Jaynes 1982). Furthermore, limiting METE to a focal taxon or taxa does not prevent accurate predictions (Harte et al. 2013), and issues of misidentification of species affect predictions in a minor way (Harte et al. 2013). We can therefore state with some certainty that tests of METE in this study are (1) robust to biases that arise in low-detectability systems, and (2) able to model the live plant community without accounting for other forms of biodiversity present in the system. These assumptions are useful to draw attention to, as they differ in important ways between the fields of macroecology and disturbance ecology. Moving towards a synthesis of the two fields will require careful study design that is informed by knowledge of both fields.

In this study, we ask how well various macroecological metrics that describe community structure, specifically the SAR, SAD, and SSADs perform at the stand level, for a forest stand that has undergone a recent (17 yr previous) disturbance in Bishop pine forests and for a nearby, mature stand in the same disturbance regime (Brown et al. 1999) at Point Reyes National Seashore (PRNS) in California, USA. We apply METE to this disturbed example systems to see if there are departures of data from theory, if the departures are systematic, and if they are coherent. We use this PRNS ecosystem as an example where we can control other covariates, but we hope to extend this analysis in future work to a broad range of biomes.

We hypothesize that the METE will predict these community structure metrics more accurately in the more mature plot (Mount Vision) because it has had a longer time since disturbance to reach steady-state dynamics, and METE’s predictions will be less accurate for the more recently disturbed (Bayview) plot.

If an information-entropy based theory of macroecology (METE) performs equally well for both the mature and disturbed plots, we would have supporting evidence that the information contained in the four state variables that constrain the predicted distributions is sufficient to describe ecosystems, regardless of their disturbance status. This would suggest that METE’s successes are independent of the disturbance history of an ecosystem. Alternately, METE might not work for one or both of the different-aged plots, which means that the theory’s four state variables do not contain adequate information to constrain the predicted distribution to the empirical distributions. Because METE is constrained to predict the maximum information entropy distributions only, the functional forms are fixed after the state variables are specified. Failures of METE to predict ecological metrics accurately in rapidly changing ecosystems would indicate the need to characterize deviations of real ecosystems from METE’s predictions, or modify the theory to allow prediction of macroecolgical metrics in ecosystems with active disturbance regimes, undergoing succession, or experiencing perturbations generally.

## Materials and Methods

### Bishop pines: a forest type that experiences natural disturbance

This study focuses on Bishop pine (*Pinus muricata*) forest stands and their associated plant communities, which exhibit an unusual natural history. Bishop pine is endemic to the California Floristic Province in North America and has a patchy distribution along the coast of California, USA and Baja California, Mexico, including the California Channel Islands (Millar 1983, Millar 1986, Little 1971, Stephens and Libby 2006) (Fig. 1). Mature stands (~40–120 years old) may have individuals that are widely spaced, and a moderately diverse understory of forbs and shrubs. Stand-replacing fires cause regeneration of the Bishop pines into a uniform age and size-class; a “dog-hair” forest that is nearly a monoculture with almost no understory. This dense forest structure been shown to undergo a process of self-thinning (Harvey et al. 2011).

**Figure 1.**
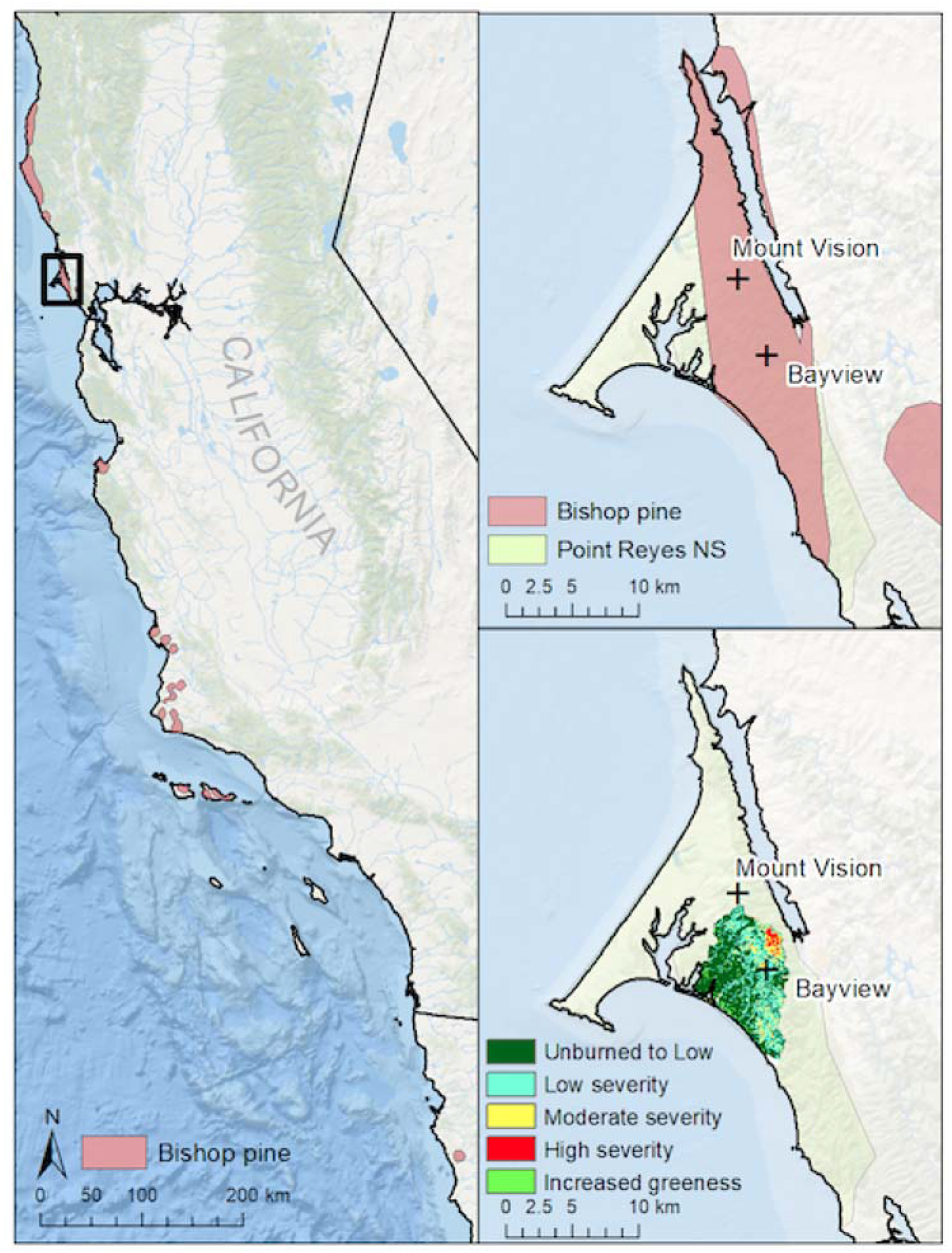
Map of Point Reyes National Seashore region in central coastal California, USA, showing study plot location and local distribution of Bishop Pines. Base maps are from Natural Earth; Bishop pine range data are from Little (1971); wildfire data from Monitoring Trends in Burn Severity (Eidenshink et al. 2007).

Alternately, some have described additional thinning fires during the lifecycle of the trees that can restore the more open canopy and allow some trees to mature into large individuals (Brown et al. 1999, S. Stephens, *pers. comm*.). Although we found no evidence for such a process at our field sites in the wildfire records maintained by Point Reyes National Seashore since the establishment of the park in 1962, Brown et al. 1999 document frequent wildfires (every 8-9 years, on average) from the 1700’s through 1945 for Olema Valley and general Point Reyes area, including 4 large fires in the early 20^th^ century (1904, 1906, 1923, and 1945). Of these, we believe the 1923 fire is the most likely to have affected our study sites. The fires are thought to be human-caused, and low-severity rather than stand-replacing. More recently, patterns of fire severity leading to landscape heterogeneity are described in Forrestel et al. (2011), which focused on vegetation succession, especially with respect to Bishop pine communities following the October 1995 Vision Fire, which burned 12,354 acres (5000 hectares, or 50 km^2^) within the National Park unit (NPS 2005). Forrestel et al. found that Bishop pines increased in extent by 85% and have an altered spatial distribution following this high-severity fire.

Species compositions between plots are not directly comparable as plant communities because the sites are exposed to different local climates. However, we are able to use our data to test hypotheses about the effects of intense natural disturbance on plant community structure from a macroecological perspective identifying overall species-level and community-level patterns, as METE’s predictions are not dependent on the identity of species in the communities.

### Site descriptions

Field sites were chosen within the boundaries of Point Reyes National Seashore (PRNS), on the Pacific coast of California, USA, ~50 km northwest of San Francisco. According to data from 1964-2012, PRNS experiences a Mediterranean-type climate, with mild winters (monthly lows of 2-4 °C and highs of 15-17 °C) and cool summers (monthly lows of 6-9 °C to highs of 18-24 °C) with most of the ~100 cm annual rainfall occurring in winter, and a substantial amount of moisture received from fog drip in the summers (Dawson 1998; Forrestel et al. 2015).

We placed study plots in two Bishop pine (*Pinus muricata*) stands, each plot measuring 256 m^2^ (16m x 16m), and censused each for all aboveground, live vascular plants ≥1 cm in height in April, 2012. Bishop pine is a serotinous species that regularly experiences stand-replacing fires. At each site, live plants were censused (all individuals counted) as completely as possible with double-observers, and each plant’s spatial location in the sampling grid was recorded with a cell number, representing a 1 m^2^ subdivision of the larger plot.

Plants were identified to species in the field when possible using the Jepson Manual (Baldwin et al. 2012) and other field guides for the local region (Howell et al. 2007, Keator and Heady 1981). In the cases where plants could not be identified to species, “morphospecies” (plants with a large number of shared characteristics) were given a unique species identifier for analysis, and reference notes and photographs were taken in the field. METE’s predictions are robust to sampling within any given taxonomic category or guild, and to the lumping or splitting of taxa, provided that such decisions are made consistently (Harte et al. 2013).

The higher elevation “Bayview” plot at 252 m (825 ft) was placed in an area of PRNS that burned in the 1995 Vision fire. We also surveyed, but discarded, a pilot plot (“Hillside”) in the 1995 Vision fire burn area because it had fewer than 10 vascular plant species and was therefore unsuitable for analysis by METE (Harte 2011; requirement that *S*_0_≫1). Both the Bayview and Hillside plots are “dog-hair” type stands of thin, closely growing trees, in which the ages of the Bishop Pines are uniform, and the understory is sparse or absent. Six trees were cored at this site to create a record of variability of widths among the center rings of growing trees following an intense fire and a period of rapid growth.

The slightly lower-elevation “Mount Vision” plot was located at 213 m (698 ft) in a mature Bishop pine stand with a more diverse and lush understory. Fourteen trees were cored at this site to estimate the ages of all trees in the plot, and results were corroborated with aerial photographs of this area in the PRNS archive (see below). The two plots, which are 6.1 km (~3.8 mi) apart are shown in Figs. 1–2, and locations and characteristics are summarized in Table 1.

**Figure 2.**
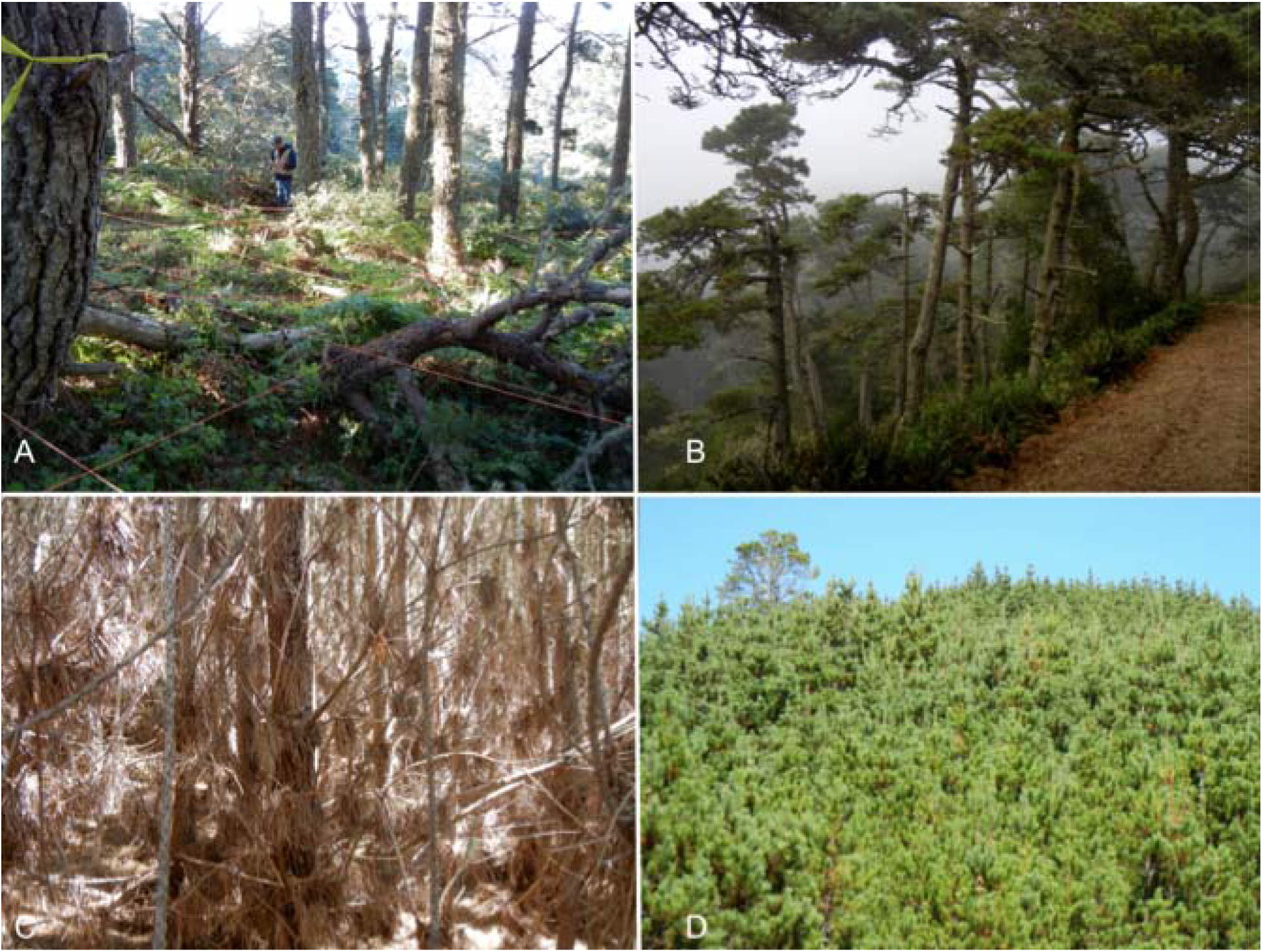
Photographs of (A) interior of mature Bishop pine stand at Mt. Vision site, and (B) side view of mature stand; (C) interior view of Bishop pine stand that burned in the 1995 Vision Fire at Bayview plot, and (D) exterior view of stand structure 17 years after the Vision Fire.

**Table 1.** Locations and other descriptive metrics for research plots used in this study

| **Site name** | **Bayview** | **Mount Vision** |
| --- | --- | --- |
| GPS coordinates<br>(lat/long) | 38.0593628°, -122.8507065° (+/- 3.6m) | 38.1028328°, -122.8933785° (+/-1.8m) |
| Elevation | 251.46 m (825 ft) | 212.75 m (698 ft) |
| Time since major disturbance | 17 years | No recorded disturbance history, although maximum tree age is measured to be 43 years |
| Slope, aspect | 0%, S | 32%, NE |
| Tree density/m<sup>2</sup> | 0.578 | 0.055 |
| Trees cored;<br>(cores available) | 4; (6) | 13; (14) |
| PRNS catalog numbers | PORE 18080 through PORE 18083;<br>PRNS Accession number: PORE-00866 | PORE 18084 through PORE 18096;<br>PRNS Accession number: PORE-00866 |
| Total species (S<sub>0</sub>) | 16 | 27 |
| Total abundance (N<sub>0</sub>) | 486 | 1844 |
| Total area (A<sub>0</sub>) | 256 m<sup>2</sup> | 256 m<sup>2</sup> |

Plant communities within Bishop pine forests at PRNS are highly patchy and exhibit high beta-diversity in the understory due to slope, elevation, and other local factors affecting climate variation, including exposure to ocean fog (Forrestel et al. 2011). As a result, species compositions between plots cannot be considered different stages of the same successional trajectory.

### Establishing disturbance histories

We examined land-use history records (including aerial photographs, contemporary accounts, historical ranch maps, and post-wildfire incident records) in the archives at Point Reyes National Seashore, in consultation with National Park Service (NPS) archival staff. Other fire records examined include CALFIRE’s Department of Forestry and Fire Protection FRAP Fire Perimeters (available online at http://frap.cdf.ca.gov/data, accessed in July 2015).

At each plot, trees were cored using increment borers (Haglöf Sweden^™^). Cores were stored in labeled paper straws until they could be glued into wooden mounts. The number of trees cored was severely limited by agreement in the National Parks permit, and a better estimate of growth height of initial growth year rings could not be obtained. See Supplementary Information: Appendix A for more information on tree ages, sampling, and curatorial information.

### The maximum information entropy approach

Census data from multiple plots within the PRNS Bishop pine community are used here to test METE predictions for the species-area relationship (SAR), the species-abundance distribution (SAD), and the species-level spatial abundance distributions (SSAD).

#### (a) Species-Area Relationship (SAR) and scale collapse

The Species-Area Relationship (SAR) describes how species diversity increases with increasing area. It is represented by 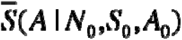, where *A* is a sampled area within the total *A*_0_ under consideration, and is calculable from the state variables as:

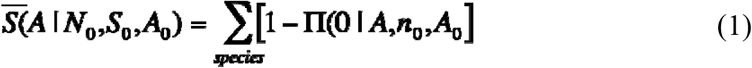

Here, Π(0 | *A,n*_0_,*A*_0_) (also known as the Species-level Spatial Abundance Distribution) is the probability that a cell (or smaller area *A* within *A*_0_) will be unoccupied by a given species, and [l-Π(0|*A*,*n*_0_,*A*_0_)] is therefore the probability of occupancy by that species. Scale collapse is a property that emerges from the METE SAR when the local slope “z” of the SAR at each spatial scale is graphed against the ratio of N/S measured at that scale (Harte et al. 2009), and we test that property for both plots compared to the METE predicted curve (Harte 2011, Harte et al. 2013, Wilber et al. 2015) with “z-D” scale collapse plots. The z-D plots show the local slope of the SAR plotted against ln(*N/S*) at every measured scale and therefore allow comparisons of SARs between plots, and even between ecosystems. The SAR calculated here is the recursive SAR, which was shown to make more accurate predictions than the non-recursive version of the same metric (McGlinn et al. 2013).

#### (b) Species-Abundance Distribution (SAD)

The SAD, **Φ**, models the distribution of the total abundance of individuals *N*_0_, across all species, *S*_0_,

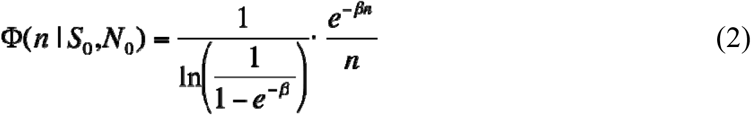

Here, *n* represents the individuals within a given species, and *β* is related to the Lagrange multipliers *λ*_1_ and *λ*_2_, which are the solutions to the information entropy maximization problem. Here, *λ*_1_= *β-λ*_2_ and *λ*_2_ = *S*_0_/(*E*_0_ – *N*_0_); and *β* satisfies the approximate relationship:

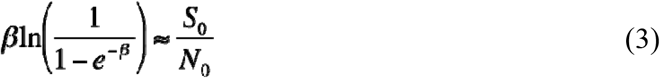

following Harte et al. 2008, Harte et al. 2009, but using exact normalization. An exact expression for *β*, its derivation, and discussion of simplifying assumptions is available in Harte (2011), Chapter 7.5. The METE SAD is here compared to the continuous lognormal distribution common to many of the studies mentioned previously, although the Poisson lognormal has also been suggested as an appropriate comparison (McGill et al. 2007, White et al. 2012).

#### (c) Species-level Spatial Abundance Distributions (SSADs)

The spatial distribution of individuals of a given species and their level of aggregation is predicted in METE as the SSAD, for which there is some support in the literature (McGlinn et al. 2015). With Π(*n* | *A*,*n*_0_,*A*_0_)(or simply Π(*n*)) defined as the species-level spatial abundance distribution, the normalization constraint on the probability distribution for a given species:

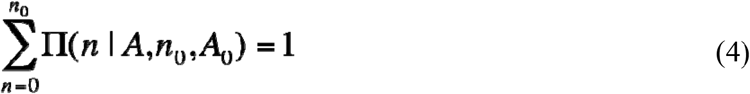

the additional constraint on the mean value of the number of individuals per cell:

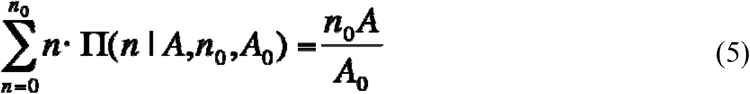

and **λ**_Π_, representing the Lagrange multiplier associated with that constraint, *Z*_Π_ is defined to be the partition function that normalizes the solutions. We can write down the form of the solution that maximizes information entropy (Jaynes 1982):

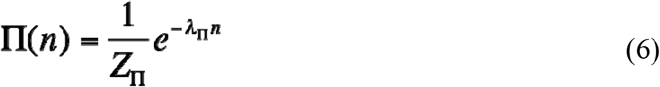

The partition function can be obtained by solving for it and the real-valued Lagrange multiplier:

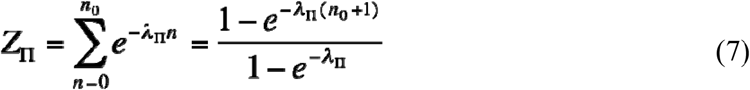

Additional steps in the solution are available in Harte (2011) Chapter 7.4.

The METE prediction for the SSAD of a given species is an “upper-truncated” (ut) geometric series, describing the frequency distribution of cells of the smallest sampled area (*A*) within the larger area *A*_0_, which contain the maximum number of individuals *n*. “Upper truncation” or “right truncation” refers to the domain of the predicted distribution being limited to its physical maximum, rather than having an unbounded upper tail of probability (see Appendix A of White et al. 2012). The METE ut-geometric prediction is here modeled with the number of parameters (*k*) of the distribution equal to 1, which is 1 parameter fewer than a regular truncated geometric. This is because the corresponding parameters in the METE distribution, *i.e*. the Lagrange multiplier and the normalization by the partition function, are uniquely determined by the same state variable values, and there are no parameters that may be adjusted in this arrangement.

### Analyses

Analyses for the macroecological metrics considered in this paper were carried out with Python (van Rossum, 2001) in the open-source project “Macroeco” (Kitzes et al. 2014, Kitzes and Wilber 2016). SAR, SAD and SSAD scripts were available from the beta version of this software (accessed June 2015). Other analyses were carried out in “R” versions 3.0.1 and 3.1.1 (R Core Team 2013, 2015). Models for SADs and SSADs were compared using Akaike’s Information Criterion (AIC) value corrected for small sample sizes (AICc). For SAR and z-D scale collapse model comparisons, we compare models with R^2^ values derived from one-to-one predicted versus observed graphs (White et al. 2012, Appendix A), because no method is available to generate likelihood functions required for AIC comparisons.

## Results

### Summary statistics and calculated parameters

A total of 2330 individual plants in 32 species were censused in the two plots analyzed for this study, though diversity and abundance differed between plots (Table 1), as did species presence (Table 2). The Bayview plot, which burned in 1995, was censused at 17 years after the Vision Fire, and contained 16 species and a total of 486 individuals (148 of which were Bishop pines). Six tree cores were aged from 8 to 16 years (mean = 13.5). Density of Bishop pines in this plot was measured to be 0.58 trees/m^2^ (or 5800 stems/ha), with a total basal area occupied by trees of 45.15 m^2^/256 m^2^ plot. The Mount Vision plot contained 27 species and 1844 individuals total (14 of which were Bishop pines). Bishop pine density in this plot measured 0.06 trees/m^2^ (or 600 stems/ha), including the very few seedling trees in the plot. The total basal area occupied by trees measured to be 17.74 m^2^/256 m^2^ plot. Tree cores varied in age from 33 to 45 years old (mean = 39.7). Live tree density per hectare estimates are consistent with estimates from other studies (Harvey et al. 2011, 2014).

**Table 2.** Presence of species by plot in two *Pinus muricata* stands of different ages. Species marked with a single asterix (*) are non-native.

| Higher taxon | Common name | Species name | Analysis code | Presence in recently disturbed (Bayview) plot | Presence in mature (Mt. Vision) plot |
| --- | --- | --- | --- | --- | --- |
| Ferns and fern allies | Sword fern | *Polystichum munitum* | POLMUN | x | x |
|  | Bracken fern | *Pteridium aquilinum* | PTEAQU | x | x |
| Gymnosperms | Bishop pine | *Pinus muricata* | PINMUR | x | x |
| Eudicots | California lilac | *Ceanothus thyrsiflorus* | CEATHY | x | — |
|  | Cape ivy* | *Delairea odorata* * | DELODO | — | x |
|  | Wood strawberry | *Fragaria vesca* | FRAVES |  | x |
|  | California coffeeberry | *Frangula californica* (syn. *Rhamnus californica*) | FRACAL | — | x |
|  | Bedstraw | *Galium sp.* (possibly *porrigens*) | GALIUM | x | x |
|  | Cow parsnip, pushki | *Heracleum maximum* | HERMAX | x | x |
|  | Cat's ear* | *Hypochaeris radicata* * | HYPRAD | — | x |
|  | Red henbit* | *Lamium purpureum* * | LAMPUR | — | x |
|  | California honeysuckle | *Lonicera hispidula* | LONHIS | x | x |
|  | False Solomon's seal, Slim Solomon | *Maianthemum stellatum* | MAISTE | — | x |
|  | California man-root, wild cucumber | *Marah fabacea* | MARFAB | x | — |
|  | Coast man-root | *Marah oregana* (syn. *Marah oreganus*) | MARORE | — | x |
|  | Wax myrtle | *Morella californica* | MORCAL | — | x |
|  | Tanoak | *Notholithocarpus densiflorus* (syn. *Lithocarpus densiflorus*) | NOTDEN | x | — |
|  | Coast live oak | *Quercus agrifolia* | QUEAGR | x | — |
|  | Canyon live oak | *Quercus chrysolepis* | QUECHR | — | x |
|  | Flowering currant | *Ribes sanguineum* | RIBSAN | x | x |
|  | California blackberry | *Rubus ursinus* | RUBURS | x | x |
|  | Sheep's sorrel, sour weed | *Rumex acetosella* * | RUMACE |  | x |
|  | Red elderberry | *Sambucus racemosa* | SAMRAC |  | x |
|  | Starflower | *Trientalis borealis* | TRIBOR |  | x |
|  | Poison oak | *Toxicodendron diversilobum* | TOXDIV | x | x |
|  | California huckleberry | *Vaccinium ovatum* | VACOVA | x | x |
|  | UNKSP3 | UNKSP3 | UNKSP3 | x |  |
|  | UNKSP5 | UNKSP5 | UNKSP5 |  | x |
| Monocots | Unknown grass | UNKPOA1 | UNKPOA1 | x | x |
|  | Unknown grass | UNKPOA2 | UNKPOA2 |  | x |
|  | Unknown grass | UNKPOA2/3 | UNKPOA3 |  | x |
|  | Unknown sedge | *Carex sp.* | CAREX |  | x |

In both plots, Bishop pines were the only tree in the overstory, and were the largest plants in each plot by estimated biomass. See Supplementary Information for histograms of dbh measurements for each site. For comparison to METE-predicted metrics, we calculate the value of the parameter *β*= 6.515 × 10^−3^ for the Bayview plot for the measured values *N*_0_ = 486, *S*_0_ = 16, and *β*= 2.429 × 10^−3^ for the Mount Vision plot, with measured values *N*_0_ = 1844, *S*_0_ = 27. The values for the Lagrange multipliers 1 and 2 are not independently calculable because state variable *E*_0_ was not measured for either plot.

### Species-Area Relationship and scale collapse

Generally, the METE prediction for the SAR appears to be a good fit for both data sets, whereas the z-D (scale collapse) predictions show more deviation from the METE prediction for the recently disturbed plot (Fig. 3). To determine the best model fits for the SAR and z-D, comparisons of *R*^2^ values on a one-to-one line for predicted versus observed distributions (White et al. 2012) were carried out for both the Bayview and Mount Vision plots. Best-fit power laws were calculated from the SAR data for each site (Fig. 3) and applied to the z-D graphs (Fig. 4). *R*^2^ values for the mature Mount Vision plot support METE’s predicted SAR over the best-fit power-law predictions (*R*^2^_METE_= 0.991; *R*^2^_powerLaw_= 0.989), whereas the power law fit is a better fit for the recently disturbed Bayview plot (*R*^2^_METE_= 0.977; *R*^2^_PowerLaw_= 0.998) on a ln-ln graph. Visually each z-D plot confirms the better fit of the METE-predicted distribution over the best-fit power-law for the mature plot.

**Figure 3.**
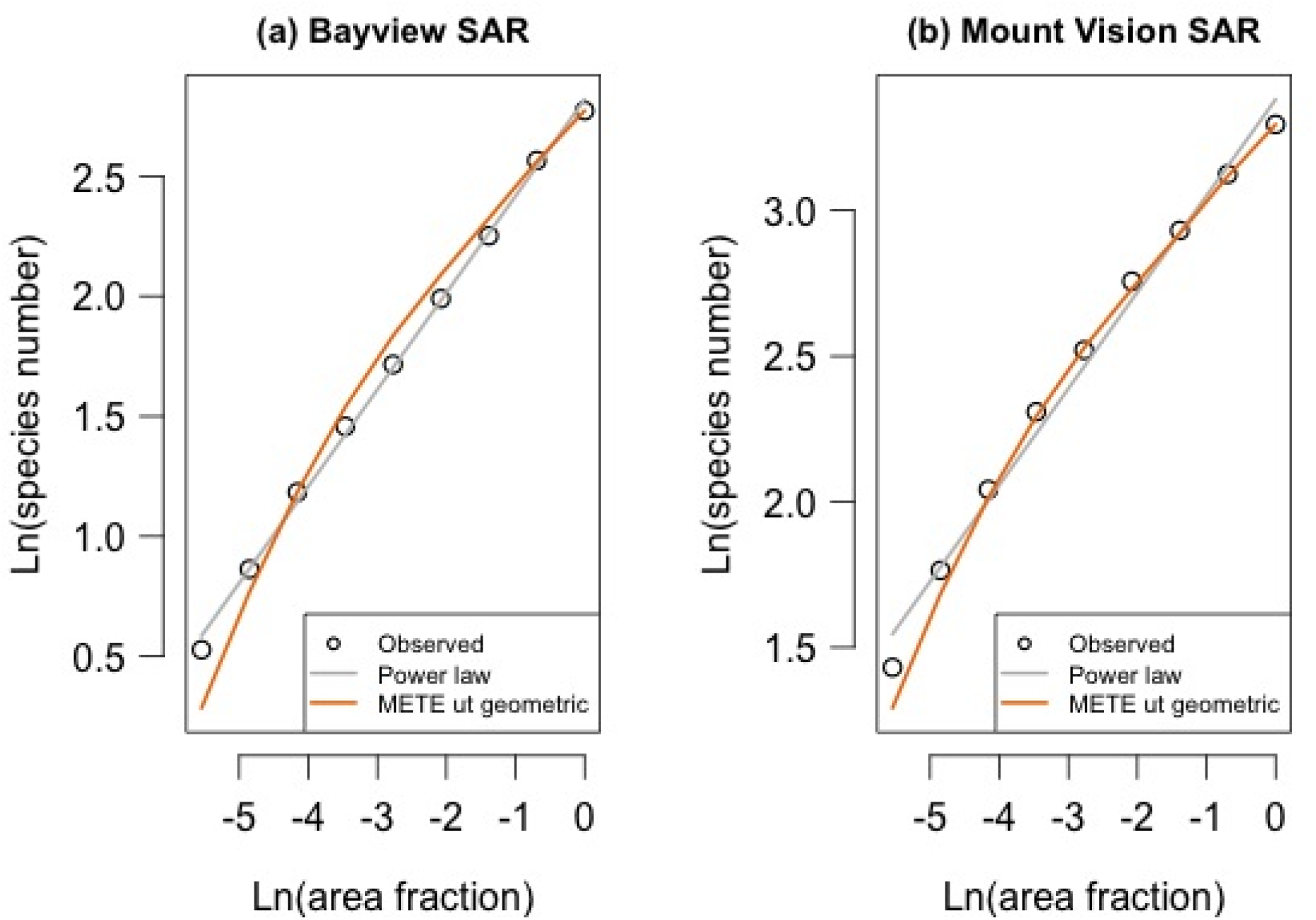
Species-Area Relationships for (a) the recently disturbed Bayview plot, and for (b) the mature Mount Vision plot. Empirical data are shown against the METE upper-truncated (ut) geometric prediction, and the best-fit power law for comparison.

**Figure 4.**
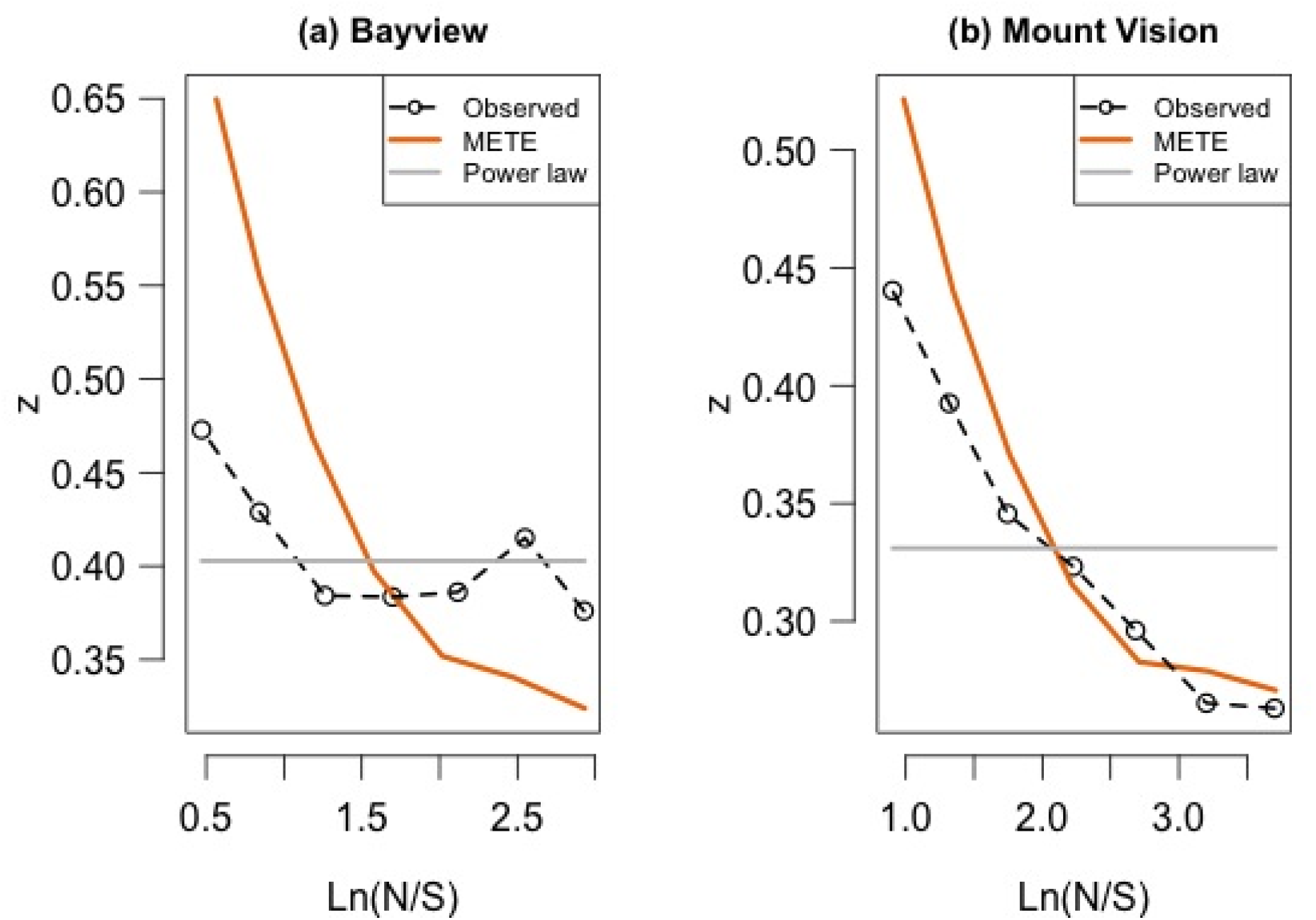
Universal scale collapse graphs, with METE-predicted and observed values illustrated at scales of N/S (total abundance/total species) for each plot. The best-fit power law is shown for comparison.

### Species-Abundance Distribution (SAD)

Model selection comparing AICc values for both the Bayview and Mount Vision plots support the METE-predicted log series distribution over the lognormal distribution (Table 3) that is sometimes characteristic of disrupted systems. On visual inspection of the SAD graphs (Fig. 5), there is the characteristic pattern of suppression of mid-abundance species in the Bayview plot that is characteristic of recently disrupted plots. The METE SAD does not capture this deviation, but because it fits the number of singleton species and the abundance of the most abundant species in this distribution, it wins out over the lognormal distribution by AICc comparisons.

**Figure 5.**
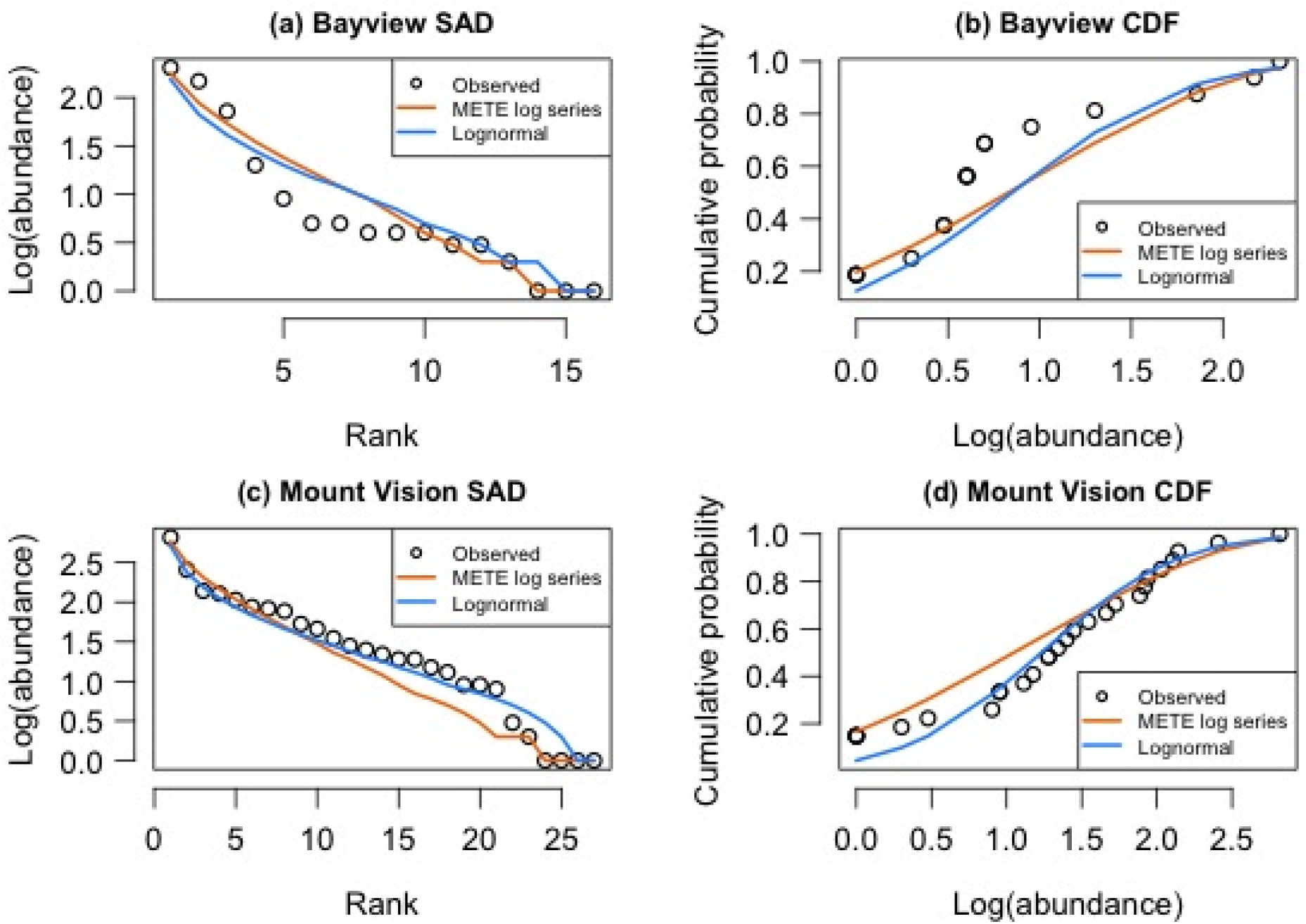
Empirical and METE-predicted ranked Species-Abundance Distributions and cumulative density functions for fire-adapted *Pinus muricata* stand in two plots: Bayview (a-b), the more recently disturbed, even-aged “dog-hair” stand that experienced a stand-replacing fire in the 1995 Vision fire, and (c-d) Mount Vision, the less recently disturbed, open stand mature trees with a more diverse understory.

**Table 3.** Model comparisons of candidate Species Abundance Distributions (SADs) for the Bayview (disturbed) and Mount Vision (mature) plots. Here and following, *k* = number of parameters in model; AICc = Akaike’s Information Criterion value corrected for small sample sizes; *w_i_* = AICc weight (a measure of strength of evidence for each model); ΔAIC*c*= difference of AIC*c* value compared to the next best-supported model.

| **Plot** | **Model** | **$k$** | **$AICc$** | **$\Delta AICc$** | **$w_i$** |
| --- | --- | --- | --- | --- | --- |
| Bayview | Lognormal | 2 | 125.3246 | 5.1485 | 0.0708 |
|  | **METE log series** | **1** | **120.1761** | **0** | **0.9292** |
| Mount Vision | Lognormal | 2 | 271.9128 | 3.7173 | 0.1349 |
|  | **METE log series** | **1** | **268.1956** | **0** | **0.8651** |

### Species-level Spatial Abundance Distributions (SSADs)

SSADs were calculated for all species. Results are presented for the higher abundance species (with *n* ≥ 20), comprising 14 species in the older Mt. Vision plot, and 4 species in the disturbed Bayview plot (Supplementary information). In Fig. 6, two alternate ways of presenting the same data are shown using the species *Trientalis borealis* (TRIBOR) as an example; first, a rank abundance plot (with rank corresponding to how many cells are occupied by a given level of abundance), and second, a cumulative density function (CDF). Note that the Poisson and binomial predictions appear to give the same results for each of the SSADs; this is because both models correspond to a null hypothesis of random placement, although the binomial has finite support and the Poisson is calculated with infinite support. AICc comparisons are summarized in Supplementary information between the candidate distributions for the SSAD: binomial, Poisson, and METE ut-geometric predictions, for the Bayview recently disturbed and Mount Vision mature plots, respectively (negative binomial fits are excluded here because they are “best-fit” and are not characterized by a single shape parameter). Supplementary information shows all CDF plots for the high-abundance species in the Bayview and Mount Vision plots, respectively. Fig. 7 shows the distribution of AICc weights for model fits for all species, for the binomial, Poisson, and METE ut-geometric predictions, with higher AICc weights corresponding to better model fits.

**Figure 6.**
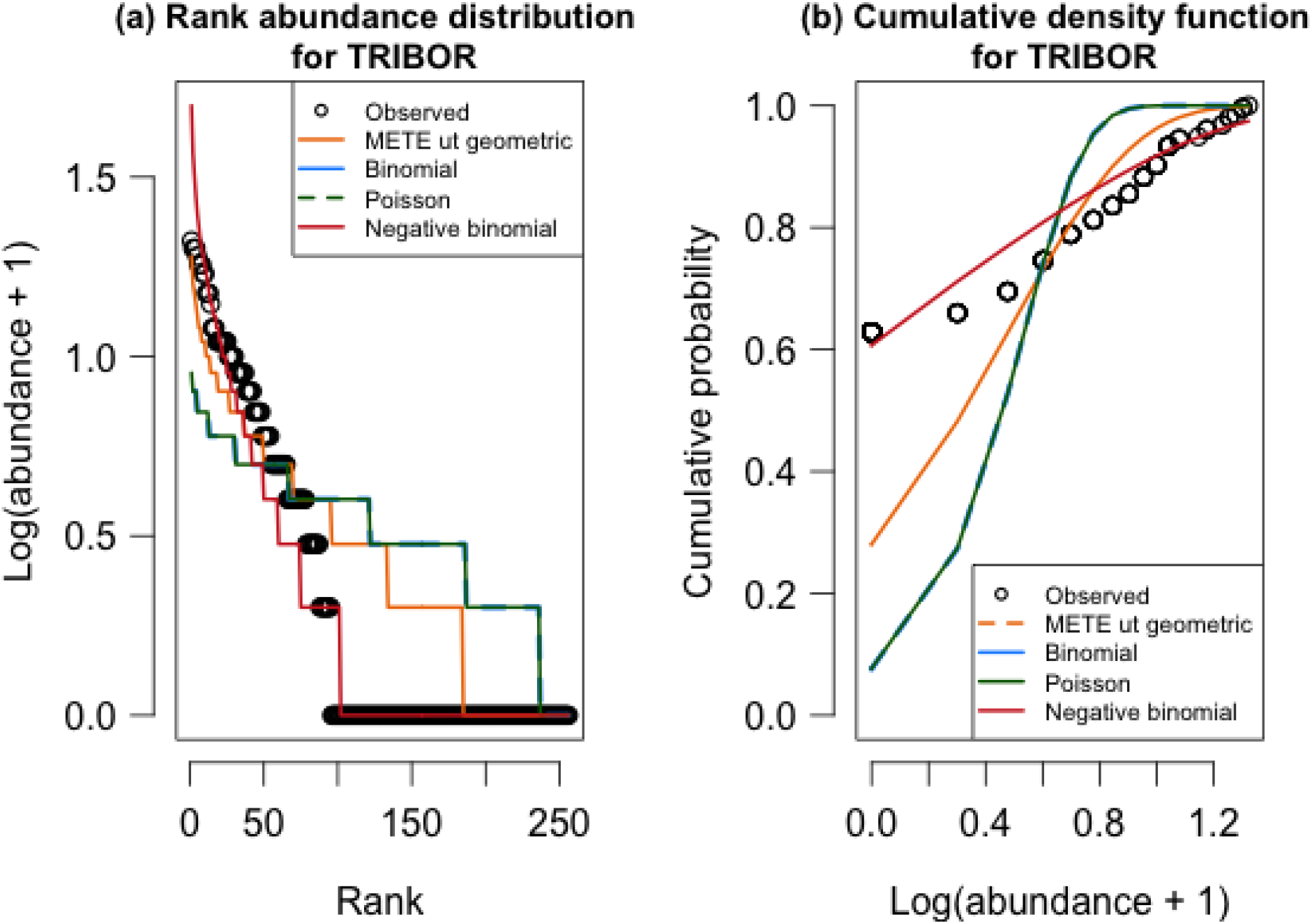
SSAD Example. Species shown is *Trientalis borealis* (TRIBOR) from the Mount Vision plot. In this example, binomial and Poisson distributions give the same predictions for the SSAD.

**Figure 7.**
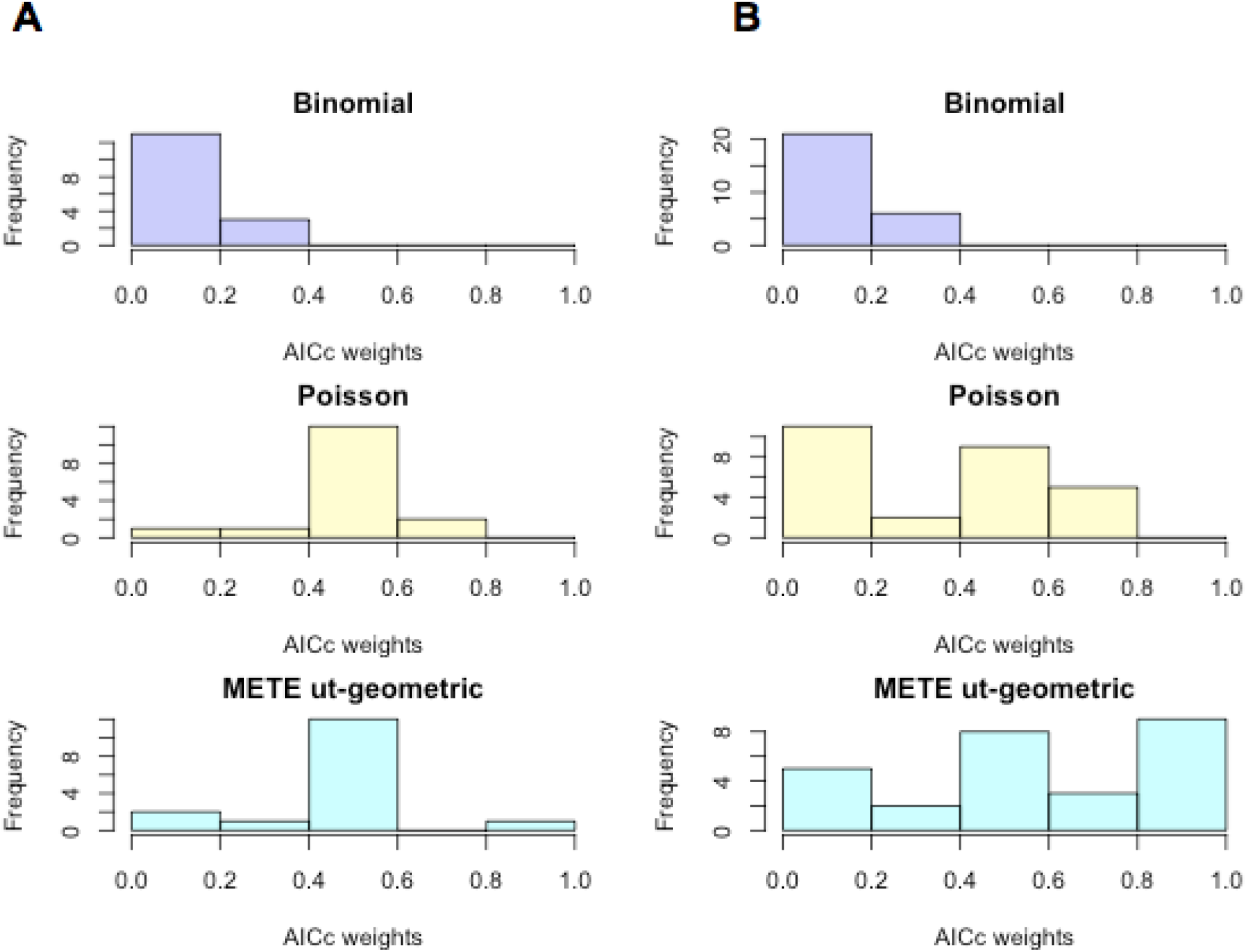
Histograms comparing SSAD models for all species by their AICc weights. By column: (A) Bayview plot (site of Vision Fire) and (B) Mount Vision mature plot. Higher AICc weights indicate better model fits.

We find that for the recently disturbed Bayview plot, SSADs for all 16 species have AICc values supporting a Poisson distribution in 11 of 16 cases, with the 5 remaining cases supporting METE ut-geometric distribution. No species are best described as having a binomial SSAD. Bishop pine distribution is best described by the METE ut-geometric distribution, with next-best supported model having ΔAICc= 3.6334. For the 16 most abundant species in the Mount Vision mature plot, AICc values support a Poisson distribution in 7 cases, and 9 cases support METE ut-geometric distribution. For all species in the plot, the Poisson distribution is the best fit for 10 species, while 17 species’ SSADs are described by the METE ut-geometric distribution (Fig. 7). Again, the distribution of Bishop pines is best described by the METE ut-geometric distribution, with next-best supported model (Poisson) having ΔAIC*c*=1.3961.

## Discussion

### Predicted and empirical distributions in different-aged stands

This study demonstrates how the field of disturbance ecology may benefit from incorporating macroecological approaches. Macroecological predictions of METE perform well in the mature stand in a disturbance-dependent community for the SAR, SAD, and the SSAD of both the dominant species (Bishop pine) and most of the other species in the plant community, compared to other candidate distributions. These results conform to our expectations, because the mature (Mount Vision) stand exhibits similar constancy and demographic stability as the extremely stable Smithsonian plots where METE has proven successful previously (Harte et al. 2008, Harte 2011, Xiao et al. 2015). METE predictions have variable and lower success in the recently disturbed (Bayview) plot.

SSAD tests in both plots would benefit from higher sample size, both in terms of number of species and individuals within species, and replicate plots. Sampling issues confounded certain results: although some two species of vines (RUBURS, TOXDIV) are among the higher abundance species at all sites, their physical description was limited to presence or absence in cells, rather than true abundance counts. As a result, the physical distribution of these species is an artifact of sampling; a high number of cells containing a single individual. This suggests that (1) for species where only occupancy can be measured, candidate models for the SSAD make degenerate predictions that can not be differentiated, therefore the SSAD is not usefully applied; (2) meaningful sampling can only be carried out at a scale where there are multiple individuals of the same species of interest in multiple cells; and (3) sampling design for tests against METE should exclude species that have this problem (resulting in changes to state variable values that will reflect only the remainder, focal community).

### Deviations from METE’s predicted distributions

METE is an effective approach for “snapshot ecology” type studies where detection rates for the taxa studied are high. However, like many forms of macroecology (Fisher et al. 2010), METE is not a dynamic theory in the ASNE formulation. Our results from two sites are consistent with the idea that ecological perturbation results in lognormal SADs, and may even be consistent with the idea that the SAD transforms during successive successional stages from a geometric shape through a lognormal to a log series (Whittaker 1975; Bazzaz 1975). It is also apparent that time since disturbance affects the shape of the SAD and various other metrics in this study, including the shift of SSADs from the Poisson towards the METE ut-geometric.

We believe deviations from METE’s predicted SAD and the more Poisson-type SSAD distributions for the general plant community in the younger stand of Bishop pines is likely explained by a lack of steady-state dynamics. As an information-entropy based statistical framework that employs state variables to describe the “macrostate” of an ecosystem or plot within that ecosystem, the static, ASNE version of METE and the MaxEnt mathematics underlying it automatically solves for the set of distributions that maximizes information entropy. This predicted state always corresponds to a steady-state solution. The fixed, maximized information entropy distributions METE predicts for a given set of state variables likely closely correspond to mature biological communities experiencing very little demographic fluctuations or other large shifts in community composition over time. This in turn may explain why METE works better for the mature stand than for the younger, more recently disturbed stand in this study. However, it still leaves open the questions of how macroecology can account for disturbance in ecosystems, and what implications this has for predicting their ecological effects.

Other examples of notable deviations from METE’s predictions have been observed in the Barro Colorado Island (BCI) forest plot, in the drought-affected Rocky Mountain (RM) meadow studied in Newman et al. 2014, and in some Hawaiian arthropod communities (Rominger et al. 2016, Harte et al. 2017). For BCI, tree and seed-disperser extirpation on the island following its isolation from the mainland was a consequence of the construction of the Panama Canal. Time since isolation has been associated with an increasingly lognormal SAD. The lognormal SAD is also observed in the RM meadow during a period of unusual drought and high temperatures leading to a novel community of wildflowers that exhibited irregular phenology (Newman et al. 2014). In the Hawai’i case, the SAD shows higher-than-predicted numbers of singleton species. Deviation from the METE in this case may be caused by dispersal limitation and the relatively young age of the community (Rominger et al. 2016). In each case, ecological context suggests that these systems are far from steady-state dynamics and provides insight into the ecological patterns observed.

### Unifying macroecology with disturbance ecology

Until this study, no macroecological studies have focused on patterns in species diversity, spatial and abundance distributions in natural disturbance regimes. This study examines two very different successional states in a disturbance dependent ecosystem in an attempt to maximize the differences between METE’s predictive abilities in communities with different disturbance histories.

We show that at the plot-scale, METE predictions are generally better supported for the more mature, less rapidly changing plot. Although METE has been demonstrated to work at the largest scale of ecosystems (Harte et al. 2009, Harte 2011, White et al. 2012, Harte and Kitzes 2015), it is unclear how well METE would predict various metrics at an intermediate “landscape scale” (~50,000-100,000 ha) that landscape contains multiple patches with different disturbance histories. Census information for this scale is generally lacking, and understanding biodiversity patterns at these large scales is part of the motivation to study macroecology (Brown 1995).

This study is a first step towards integrating macroecology into the study of landscapes undergoing natural disturbances, but examining single successional states will not capture integral aspects of disturbance-dependent ecosystems. For example, the dynamics of the “shifting mosaic” of different successional states and patches that themselves may exhibit dynamic steady-states with respect to regional climate (Bormann and Likens 1979, Wu and Loucks 1995). As a form of “snapshot ecology” that incorporates no dynamics, it is possible that the ASNE version of METE may best be applied to patches within a disturbed landscape to characterize zones of different ages and disturbance histories. Alternately, measuring state variables from multiple patches at different successional stages will provide more constraining information for METE metrics, and may lead to better predictions of the SAR and SAD. The scaling of macroecological metrics may even provide insight into the scales at which disturbance-dependent ecosystems deviate most dramatically from METE predictions, as patterns in disturbance-dependent landscapes are known to be scale-dependent (Wu 2004). These patterns may indeed change between landscapes characterized by large, infrequent disturbances (as in this study) and small, frequent disturbances (Romme et al. 1998, Turner and Dale 1998).

### Implications for conservation of Bishop pine forests

Although these Bishop pine forests provide a window onto the macroecology of natural disturbances, anthropogenic disruptions have become dominant in this ecosystem. Ongoing infestation by a non-native pathogen (Pitch Canker, *Fusarium circinatum*) is rapidly causing Bishop pine mortality over large, continuous areas within the Vision Fire burned area. Mortality in some places is close to 100% (Ben Becker, *pers. comm*.). This pathogen will likely permanently affect stand structure and viability of Bishop pines, and may endanger them as a species. Understanding stand structure and heterogeneity in unaffected stands while they are still present may be critical to conservation efforts.

